# Microtubule-mitochondrial attachment facilitates cell division symmetry and proper mitochondrial partitioning in fission yeast

**DOI:** 10.1101/2021.12.19.473396

**Authors:** Leeba Ann Chacko, Felix Mikus, Nicholas Ariotti, Gautam Dey, Vaishnavi Ananthanarayanan

## Abstract

Association with microtubules inhibits the fission of mitochondria in *Schizosachharomyces pombe*. Here we show that this attachment of mitochondria to microtubules is an important cell intrinsic factor in determining division symmetry. By comparing mutant cells that exhibited enhanced attachment and no attachment of mitochondria to microtubules (Dnm1Δ and Mmb1Δ respectively), we show that microtubules in these mutants displayed aberrant dynamics compared to wild-type cells, which resulted in errors in nuclear positioning. This translated to cell division asymmetry in a significant proportion of both Dnm1Δ and Mmb1Δ cells. Asymmetric division in Dnm1Δ and Mmb1Δ cells resulted in unequal distribution of mitochondria, with the daughter cell that received more mitochondria growing faster than the other daughter. Taken together, we show the existence of homeostatic feedback controls between mitochondria and microtubules in fission yeast, which directly influence mitochondrial partitioning and thereby, cell growth.

## Introduction

Symmetric cell division is the hallmark of most eukaryotic cells. Fission yeast (*Schizosaccharomyces pombe*) is a rodshaped, unicellular eukaryote that divides symmetrically during mitosis (1). A single cell grows by polarised tip extension from about 7μm to 14μm in length. Once the cell has grown to 14μm in length, it ceases to grow and proceeds to divide by assembling an actomyosin contractile ring at the geometrical centre of the cell (2, 3). Subsequently, the two daughter cells formed post mitosis are of equal length. Due to their ability to divide medially and produce identically-sized daughter cells, fission yeast is a powerful tool in cell cycle research.

One of the key players involved in ensuring symmetric division in fission yeast has been identified to be the microtubule (MT) cytoskeleton (4). A typical fission yeast cell contains an average of three to five MT bundles that emanate in the perinuclear region from the centrosome (spindle pole body in yeast) or other interphase MT organising centres (iMTOCs) (5), and are positioned along the long axis of the cell (6). MTs in *S. pombe* can crossbridge with the nuclear envelope (6), and iMTOCs themselves are thought to interact with the nuclear envelope (4). The pushing forces of the individual bundles growing against the cell periphery in an interphase cell ensure the medial placement of the nucleus (4). This medial placement enables positioning of the division plane at the centre of the cell (7). As a result, attenuating the dynamics of MTs causes severe cell division defects.

Contrary to their depiction in textbooks, mitochondria are not discrete, static entities, but rather a network of tubules that are in an equilibrium between fission and fusion. This balance between fission and fusion is essential for proper mitochondrial function, with dysfunction being associated with several cellular metabolic defects (8). The dynamin-related GTPase Drp1 (Dnm1 in yeast) is the major mitochondrial fission protein, whereas two sets of GTPases Mfn1/2 and Opa1 bring about fusion of the outer membrane and inner membrane of the mitochondria respectively (9–11). Dnm1 is cytosolic but assembles as rings around the mitochondrial outer membrane and undergoes GTP hydrolysis to effect constriction and eventual scission of mitochondria (12, 13). In the absence of Dnm1, mitochondria exist as a single, long network that spans the entire cell, but remains attached to the MT (14).

In fission yeast, mitochondria are bound to MTs via the linker protein Mmb1 (15). Recently, we showed that the absence of Mmb1 results in mitochondrial fragmentation due to the inability of Dnm1 to assemble around mitochondria bound to MTs (16). In cells with shorter MTs than normal, we observed several shorter mitochondria, whereas in cells with longer MTs than wild-type (WT), we observed fewer, longer mitochondria. Importantly, the total mitochondrial volume between the WT cells and mutant strains with shorter or longer MTs was conserved, indicating that the predominant result of altered MT dynamics was a change in mitochondrial morphology. We therefore established a causal link between MT dynamics and mitochondrial morphology (16).

In this work, we explore the outcome of altered mitochondrial form, and thereby their attachment to MTs in context of cell division. We observed that both Dnm1Δ and Mmb1Δ cells displayed increased asymmetric cell division. We set out to investigate the mechanism by which alteration of mitochondrial form resulted in these cellular homeostasis defects.

## Results

### Dnm1Δ and Mmb1Δ exhibit asymmetry during cell division

Cells lacking the mitochondrial fission protein Dnm1 contain a single long mitochondrial network ((14), Fig S1A). This long mitochondrion was attached to MTs along the length of the cell, such that when MTs were depolymerised using MBC (methyl-2-benzimidazole-carbamate), we observed retraction of the mitochondrial network (Fig. S1B, Video S1). This evinced that there was an enhanced attachment of mitochondria to MTs in Dnm1Δ cells. On the other hand, cells lacking the mitochondria-MT linker protein Mmb1 do not associate with MTs (15). We confirmed these observations by visualising MTs and mitochondria in ultrastructure-expanded images (17) of WT, Dnm1Δ and Mmb1Δ cells (1A, Video S2), and indeed quantified higher rates of attachment of mitochondria to MTs in Dnm1Δ cells compared to WT cells (1B). In our previous work, we showed that this dissociation of mitochondria from MTs results in fragmentation of the mitochondrial network (Fig. S1A, (16)). When we followed dividing Dnm1Δ and Mmb1Δ cells, we observed that cells exhibited ∼15% asymmetry in both cell length and cell area during division, compared to a median of ∼5% asymmetry in WT cells (Fig. 1C-E, S1C). Accordingly, the daughter cells in Dnm1Δ and Mmb1Δ background were also distributed across a larger range of areas than the WT cells (Fig. 1F). This degree of asymmetry during division is significantly higher than WT cells but slightly lower compared to the phenotype in Klp4Δ (MT-stabilising kinesin-like protein (18), Fig. 1), Pom1Δ (polarity-determining protein kinase (19), Fig. S1D, E) which have been well-established to exhibit asymmetry in division. Cells lacking the heteromeric kinesin-8 Klp5/6 have longer MTs and mitochondria than WT (16), and therefore also have increased attachment of mitochondria to MTs. Klp5/6Δ cells also showed increased asymmetric division compared to WT cells (Fig. S1D, E).

**Fig. 1.**
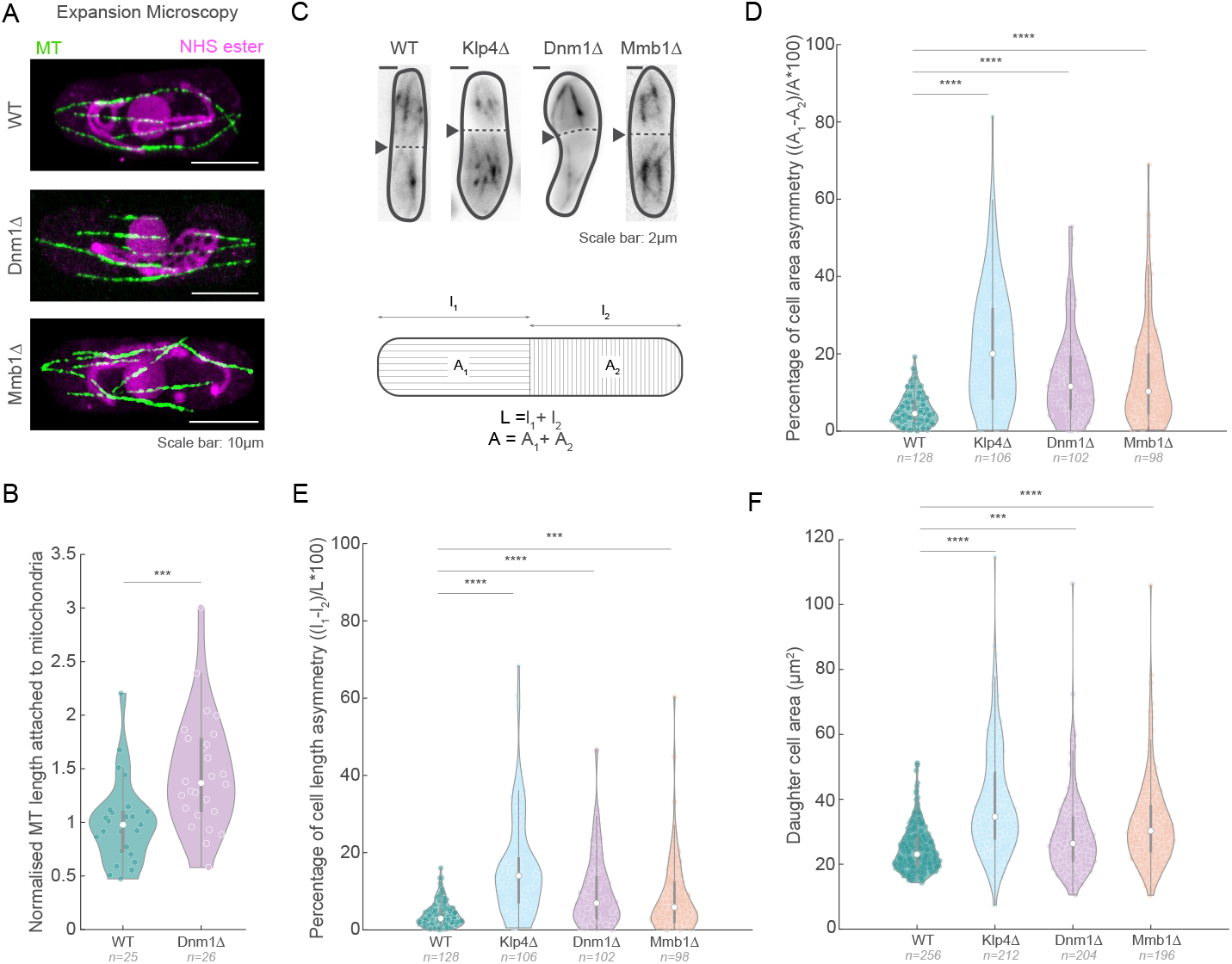
Dnm1Δ and Mmb1Δ cells exhibit increased asymmetric cell division. **A**, Spinning disk confocal microscopy images of MTs (green) and NHS ester (magenta) in ultrastructure-expanded WT, Dnm1Δ and Mmb1Δ cells (strains L972, Dnm1Δ and VA078, see Table S1). The NHS ester non-specifically labels protein density, particularly the mitochondria and nucleus, as seen in these cells. **B**, Quantification of MT length attached to mitochondria in WT and Dnm1Δ cells normalised to the mean of WT cells. Note that Mmb1Δ cells do not show attachment between MTs and mitochondria. **C**, Maximum intensity projected images of MTs in WT, Klp4Δ, Dnm1Δ and Mmb1Δ (strains L972, FY7143, KI001, G5B, Dnm1Δ and VA069, see Table S1), with the cell division plane (dashed line) indicated with the black arrowheads (top), and schematic of the method employed to measure cell length and area asymmetries (bottom). **D**, Plot of asymmetry in cell areas between the daughter cells in WT, Klp4Δ, Dnm1Δ and Mmb1Δ cells. **E**, Plot of asymmetry in cell lengths between the daughter cells in WT, Klp4Δ, Dnm1Δ and Mmb1Δ cells. **F**, Plot of daughter cell area in WT, Klp4Δ, Dnm1Δ and Mmb1Δ cells. In **B**, the asterisks represent significance (*** = p<10^−3^, student’s T-test). In **D, E** and **F**, the asterisks represent significance (**** = p<10^−4^ and *** = p<2×10^−4^ respectively), Kruskal-Wallis test for non-parametric data.

We asked if the asymmetry could have arisen due to defects in mitochondrial function in the mutant cells. To answer this question, we quantified the proportion of asymmetry in dividing *rho*^0^ cells. *S. pombe* cells are petite negative and as such, these *rho*^0^ cells have an additional nuclear mutation to grow in the absence of mtDNA (20, 21). These *rho*^0^ cells rely primarily on glycolysis for ATP production, and therefore grow slower on fermentable carbon sources (20). We did not observe significant differences in cell division asymmetry between WT and *rho*^0^ cells (Fig. S1D, E). Mitochondrial form is also linked to reactive oxygen species (ROS) levels, with fragmented mitochondria producing increased ROS and fused mitochondria producing reduced ROS (22). However, from previous work, we did not see a difference in mitochondrial ROS in mutants with altered mitochondrial morphology (16). So too, transformation of Dnm1Δ cells with Dnm1 restored mitochondrial form (16) and also symmetry in daughter cell length during division (Fig. S1C, D).

### Microtubule dynamics are altered in mitochondrial morphology mutants

Nuclear positioning in *S. pombe* is effected by pushing forces of growing MTs against the cell poles (4). Due to the paired anti-parallel nature of MT bundles in fission yeast (5, 6), this translates to net equal forces on either side of the cell. Therefore, the nucleus largely remains in the centre of the cell and this central location of the nucleus is essential in dictating the future cell division plane. Fission yeast MT mutants, such as Klp4Δ and Klp5/6Δ, have altered MT dynamics, and therefore mis-center the nucleus, leading to a significant increase in asymmetrically dividing cells (Figs. 1C-E, S1C-E). We asked whether Dnm1Δ and Mmb1Δ cells displayed asymmetry in cell division due to altered MT dynamics. Mmb1Δ cells have been described to have more dynamic MTs than WT cells, and cells over-expressing Mmb1 exhibit more stable MTs (15). So too, Dnm1Δ cells required a higher concentration of the MT-depolymerising drug TBZ to completely abrogate MTs (14), indicating higher MT stability. We measured the MT polymerisation rate, depolymerisation rate and MT elongation time in WT, Klp4Δ, Dnm1Δ and Mmb1Δ cells (Fig. 2A), and observed that MTs in Dnm1Δ cells had reduced depolymerisation rate (Fig. 2C) and increased elongation time (reduced catastrophe frequency) compared to WT cells (Fig. 2D). On the other hand, Mmb1Δ cells had MTs with increased depolymerisation rate (Fig. 2C). As expected, Klp4Δ cells exhibited reduced MT depolymerisation rate and polymerisation rate compared to WT cells (Fig. 2A, B and C). These results indicated that the association of mitochondria with MTs enhanced MT stability, whereas the lack of association reduced MT stability. We confirmed that these results were not an artefact of the levels of tubulin expression in these cells by comparing the total intensity of tubulin among the strains employed (Fig S2A).

**Fig. 2.**
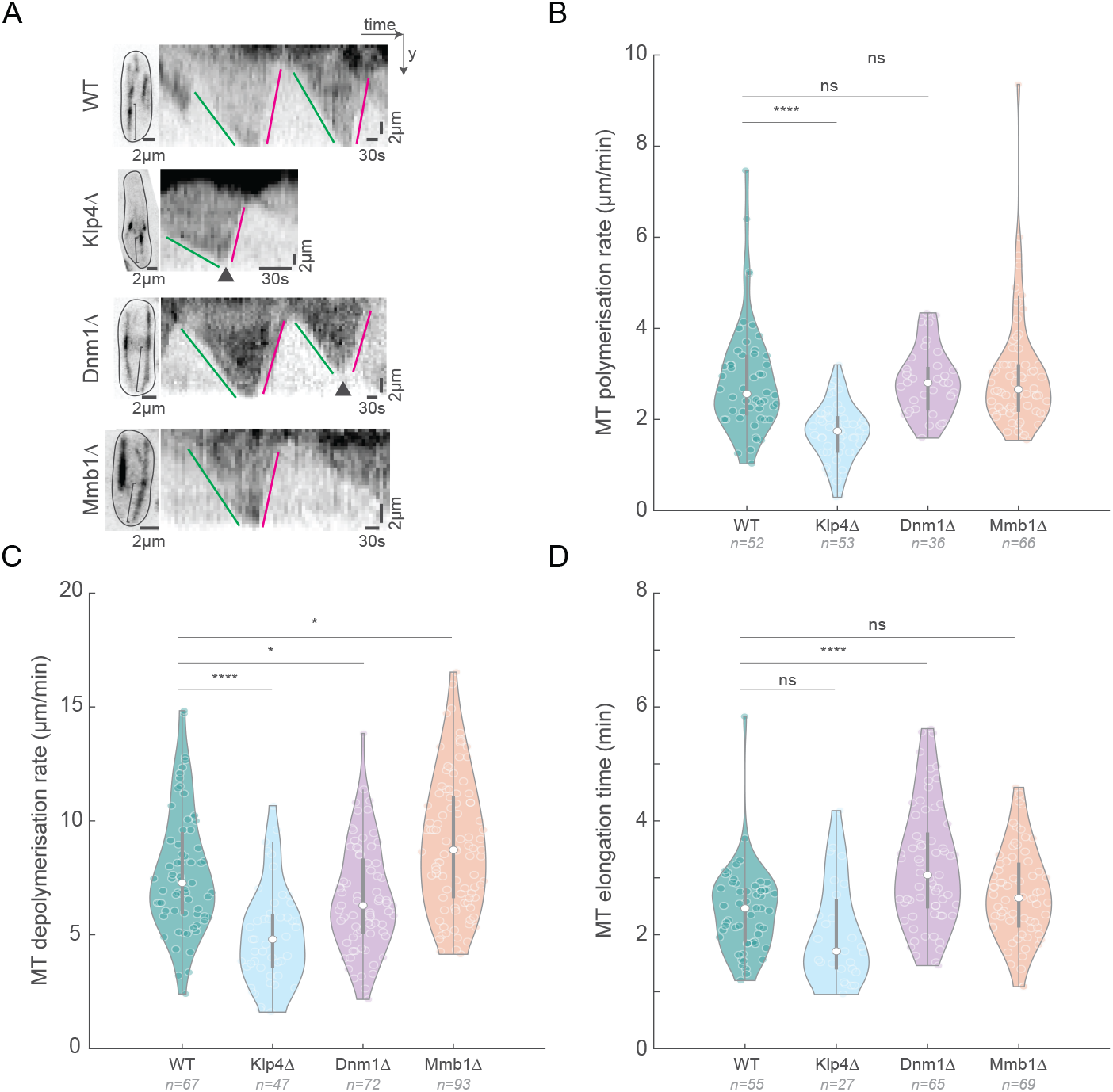
MT depolymerisation rate is aberrant in Dnm1Δ and Mmb1Δ cells. **A**, Maximum intensity-projected images (left) of MTs from the first frame of time-lapse videos of representative WT, Klp4Δ, Dnm1Δ and Mmb1Δ cells (strains VA112, G5B, VA110 and VA113, see Table S1), and the corresponding kymographs (right) of the MTs indicated with the square brace. Green lines indicate MT polymerisation, magenta lines indicate MT depolymerisation and the arrowheads point to catastrophe events. **B**, Plot of MT polymerisation rates in WT, Klp4Δ, Dnm1Δ and Mmb1Δ cells (mean *±* S.D.: 2.9 *±* 1.2, 1.7 *±* 0.6, 2.8197 *±* 0.7, and 3.0 *±* 1.3 *μ*m/min respectively). **C**, Plot of MT depolymerisation rates in WT, Klp4Δ, Dnm1Δ and Mmb1Δ cells (mean *±* S.D.: 7.8 *±* 2.7, 5.0 *±* 2.1, 6.7 *±* 2.3, and 9.0 *±* 2.9 *μ*m/min respectively). **D**, Plot of MT elongation times in WT, Klp4Δ, Dnm1Δ and Mmb1Δ cells (mean *±* S.D.: 2.4 *±* 0.7, 2.0 *±* 0.9, 3.2 *±* 1.1, and 2.7 *±* 0.8 min respectively). The reciprocal of the MT elongation time gives the MT catastrophe rate. In **B, C** and **D**, the asterisks represent significance (**** = p<10 and * = p<11×10 respectively), and ‘ns’ indicates no significant difference using Kruskal-Wallis test for non-parametric data and ordinary one-way ANOVA for parametric data.

### The nucleus is highly dynamic in mitochondrial morphology mutants

Since the nuclear position prior to onset of mitosis determines the future site of division (4), we set out to ask if the altered MT dynamics in the mitochondrial morphology mutants changed the nuclear dynamics in these cells. We observed that unlike WT cells, the nucleus was highly dynamic in both Dnm1Δ and Mmb1Δ cells (Fig. 3A, Video S3). As a result, the excursions of the nucleus from the cell centre were significantly higher in Dnm1Δ and Mmb1Δ cells than in WT cells (Fig. 3B). We confirmed that the nucleus moved more as a result of the altered MT dynamics by visualising the nuclear dynamics in cells devoid of MT (Fig. S2B). As expected, we measured negligible movement of the nucleus in the absence of MTs. So too, the short MTs in Klp4Δ cells typically do not contact the cell end (16, 18), and therefore does not result in a pushing force to move the nucleus. This was reflected in the reduced movement of the nucleus (Fig. 3A, Video S3), and increased distance of the Klp4Δ nuclei from the cell centre (Fig. 3B). Occasionally, we observed Dnm1Δ and Mmb1Δ cells that had inherited few or no mitochondria from the mother cell. Remarkably, the nuclei in these cells exhibited dramatic movements, reiterating that MT instability could be effected by lack of mitochondrial attachment (Fig. S2C, D, Video S4).

**Fig. 3.**
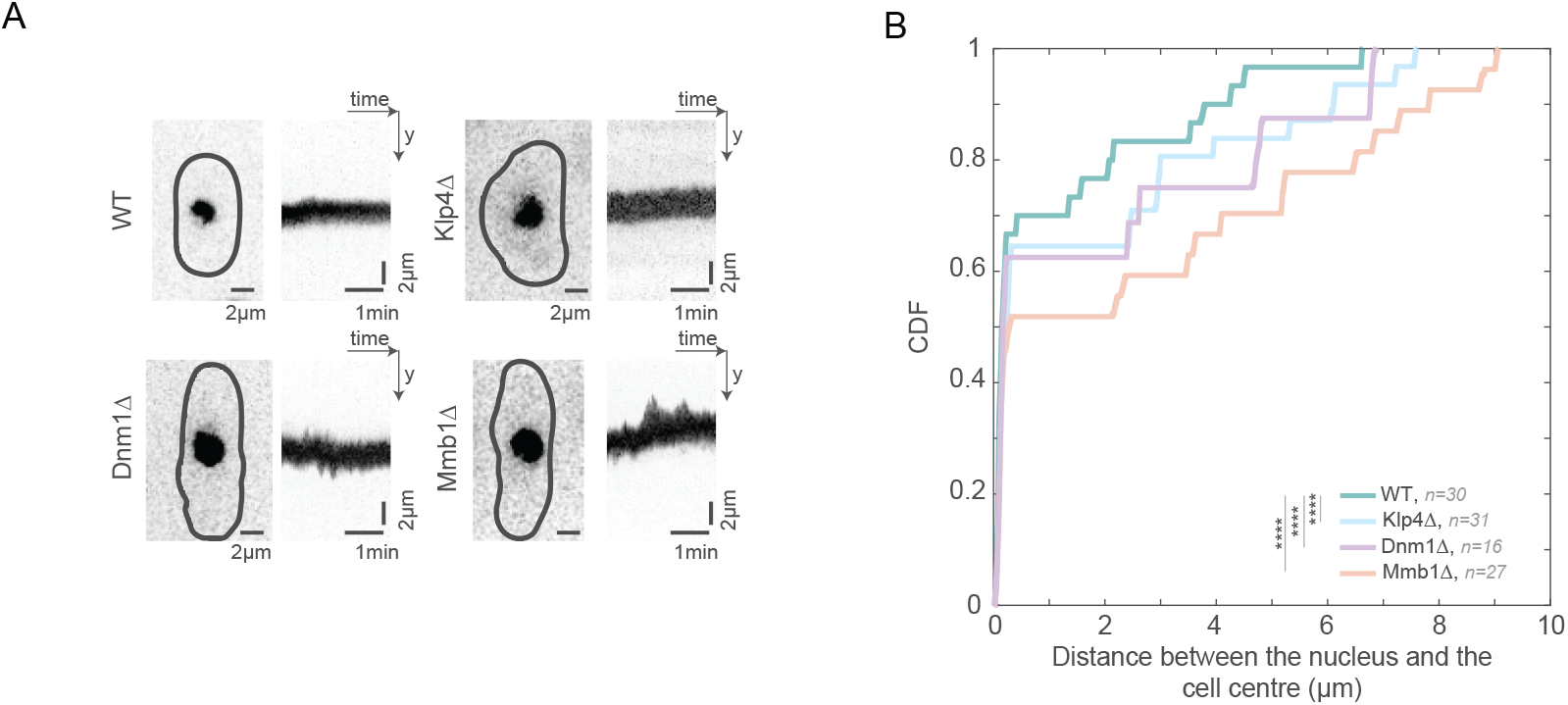
Dnm1Δ and Mmb1Δ cells exhibit enhanced nuclear movement. **A**, Maximum intensity-projected images (left) of the nucleus from the first frame of time-lapse videos of representative WT, Klp4Δ, Dnm1Δ and Mmb1Δ cells (strains VA102, VA111, VA103 and VA104, see Table S1), and the corresponding kymographs (right) of the nuclear movement. **B**, Cumulative density function (CDF) of the distance of the nucleus from the cell centre for each time point of the time-lapse videos of nuclei in WT, Klp4Δ, Dnm1Δ and Mmb1Δ cells. The asterisks (****) represent p<10^−4^, Kruskal-Wallis test for non-parametric data.

### Mitochondrial partitioning is asymmetric in mitochondrial morphology mutants

Next, we probed the consequence of asymmetric division of mutant cells on the partitioning of mitochondria. Mitochondria undergo independent segregation in fission yeast, with cell division symmetry aiding the equitable partitioning of mitochondria between daughter cells (16). We measured the amount of mitochondria in dividing WT and mutant cells (Fig. 4A), and observed that mitochondria were partitioned in proportion to the cell area, indicating that independent segregation was still likely active in the mutants (Fig. 4B). However, since a significant proportion of cells underwent asymmetric division in the mutants, mitochondria were also partitioned unequally between daughter cells (Fig. 4C).

**Fig. 4.**
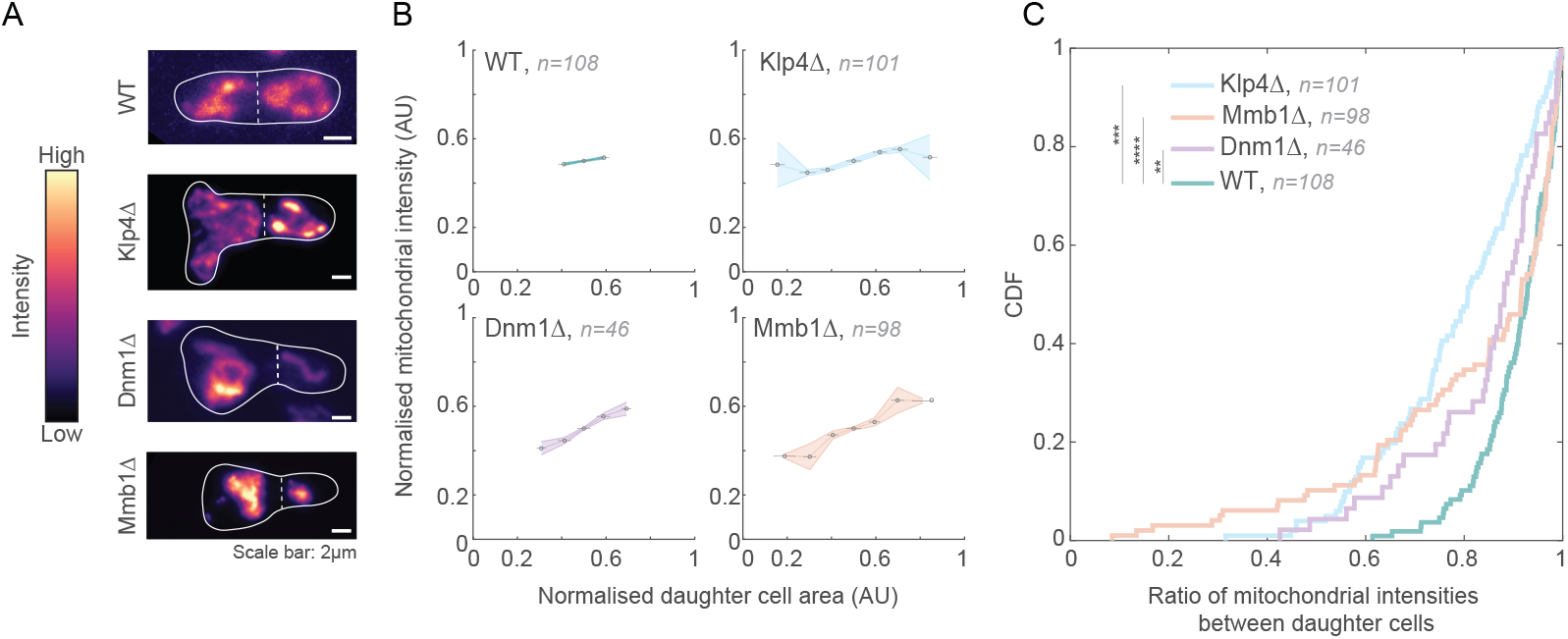
Mitochondria are asymmetrically partitioned in Dnm1Δ and Mmb1Δ cells. **A**, Maximum intensity-projected images of mitochondria in WT, Klp4Δ, Dnm1Δ and Mmb1Δ cells (strains KI001, G5B, VA069 and PT2244, see Table S1). Warmer colours indicate higher intensities. The cell outlines are indicated with the solid white line and the septum between the daughter cells is the dashed white line. **B**, Plots of normalised mitochondrial intensity (sum intensity) vs. normalised cell area in WT, Klp4Δ, Dnm1Δ and Mmb1Δ cells. **C**, CDF of ratio of mitochondrial intensities between daughter cells in WT, Klp4Δ, Dnm1Δ and Mmb1Δ cells. The asterisks represent significance (**** = p<10^−21^, *** = p<10^−10^ and ** = p<3×10^−6^ respectively) using Levene’s Test for equality of variances.

### Growth rate of cells scales with quantity of mitochondria inherited following cell division

Finally, we tested the outcome of such asymmetric partitioning of mitochondria in Dnm1Δ cells that underwent asymmetric cell division. We observed that the smaller daughter, which received less mitochondria than the larger daughter, grew slower than the larger daughter cell (Fig. 5A, B, S2E, Video S5). In comparison, WT cells which show only a small degree of asymmetry in cell area (∼5% on average) and therefore mitochondrial partitioning, still exhibited differences in growth rates between the two daughters (Fig. 5B).

**Fig. 5.**
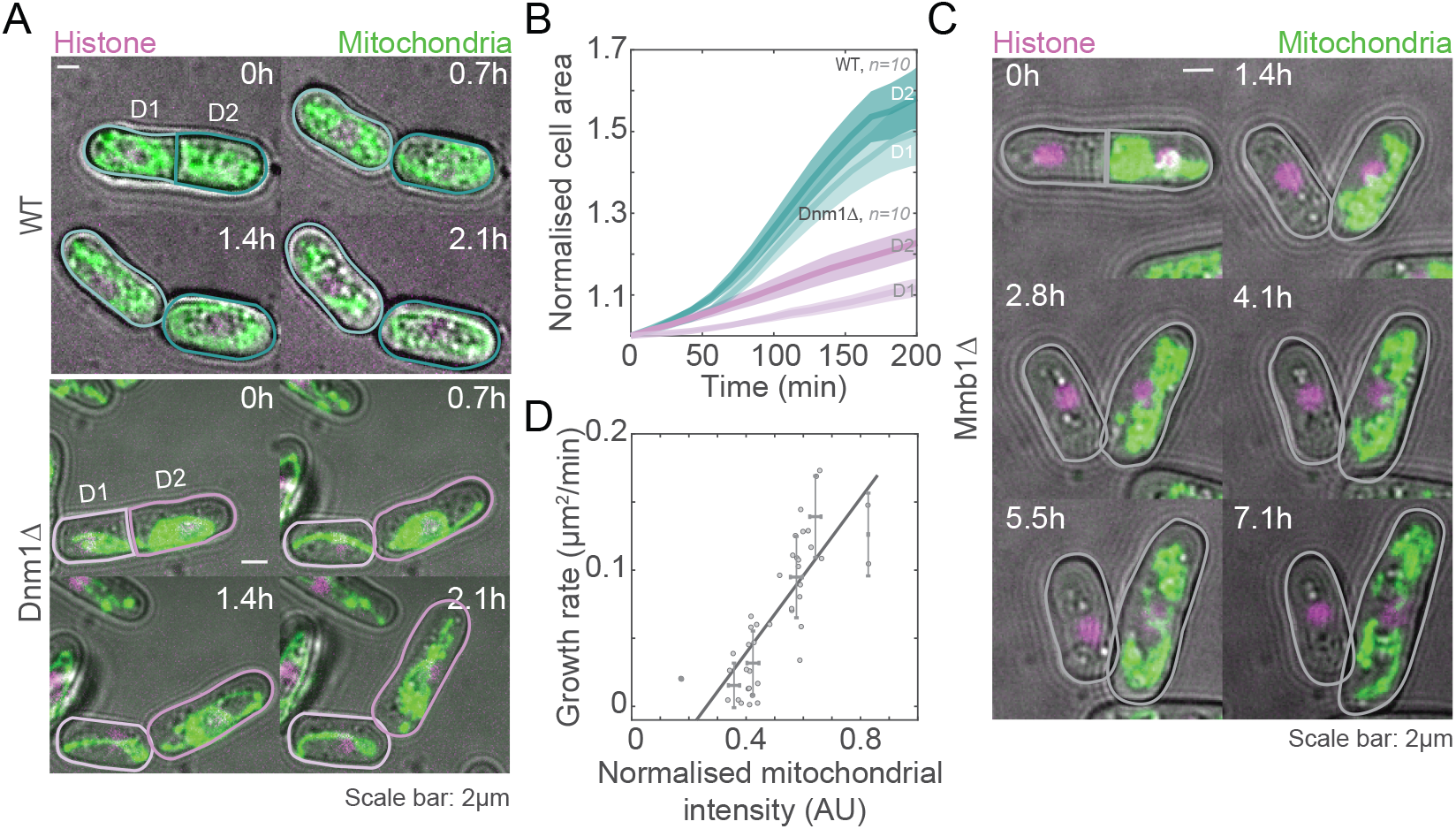
Mitochondrial content at cell birth determines growth rate. **A**, Montage of maximum intensity-projected images of mitochondria (green) and histone (magenta) in a representative WT cell (top, strain VA102, see Table S1) and Dnm1Δ cell (bottom, strain VA103, see Table S1) undergoing asymmetric cell division and mitochondrial partitioning. D1 is the smaller daughter cell, and D2, the larger daughter cell. **B**, Plot of change in cell area of D1 and D2 cells vs. time normalised to the first time frame upon division of the cells. 10 D1-D2 pairs were analysed for WT and Dnm1Δ cells (strains VA102 and VA103, see Table S1). **C**, Montage of maximum intensity-projected images of mitochondria (green) and histone (magenta) in a representative Mmb1Δ cell (strain VA104, see Table S1) with symmetric cell division but asymmetric mitochondrial partitioning. **D**, Plot of growth rate vs. mitochondrial intensities in 21 Mmb1Δ daughter cell pairs that underwent <20% asymmetric cell division. The black line is a weighted linear fit (of the form *y* = *mx* + *c*), and yielded *R*^2^ =0.81.

We confirmed that the growth rates were proportional to the mitochondria inherited from the mother by quantifying the growth rates in symmetrically dividing cells that partitioned mitochondria asymmetrically. Such events are occasionally seen in Mmb1Δ cells (Fig. 5C, D, S2F, Video S6). We observed a linear relationship between mitochondrial inheritance at the time of birth and the growth rate (Fig. 5D), indicating a central role for mitochondria in determining dynamics of cell growth.

## Discussion

The interplay between mitochondria and MTs has been implicated in maintaining cellular homeostasis. Here, we first identified that alteration of mitochondrial form and thereby, attachment of mitochondria to MTs resulted in higher rates of incidence of asymmetry in typically symmetrically-dividing fission yeast cells. We showed that this asymmetry resulted from changes in MT depolymerisation rate and catastrophe frequency when the association of mitochondria to MTs was either enhanced or absent compared to WT cells. In metazoans, mitochondria rely on microtubules for their transport and positioning (23). Further, MTs in metazoans have been demonstrated to effect changes in gene expression owing to their link with the nuclear membrane via the LINC (linker of nucleoskeleton and cytoskeleton) complex (24). It would be interesting to see if a change in mitochondrial form or attachment to MTs has a similar effect on MT dynamics, and therefore cell fate in metazoans.

The endoplasmic reticulum (ER), another prominent organelle in most cells, has been recently shown to have a mechanical role in controlling MT organisation in mammalian cells (25), and in constraining spindle lengths in *Drosophila* syncytial embryos (26), providing additional evidence for general organelle-mediated MT regulation. In *S*.*pombe*, the ER is not known to directly associate with MTs. However, there may be indirect links between these two components via the ER mitochondria encounter structures (ER-MES), which regulate mitochondrial form and biogenesis (27).

The perturbation of MT dynamics in fission yeast mutants with altered mitochondrial form resulted in increased nuclear movements, which gave rise to nuclear positioning that was offset from the cell centre. Since fission yeast relies on the nuclear position prior to mitosis to dictate the eventual cell division plane, mutants with altered mitochondrial form exhibited more instances of asymmetric cell division compared to WT cells.

Fission yeast as well as other metazoans have been documented to follow independent segregation to partition mitochondria among daughter cells during mitosis (16, 28). Independent segregation relies on the presence of a large ‘copy number’ of mitochondria present in the mother cell so as to reduce the partitioning error (29). Given large enough copy numbers of mitochondria, positioning the division plane roughly at the cell centre ensures equitable distribution of mitochondria in daughter cells. In Mmb1Δ and Dnm1Δ cells, due to the asymmetry observed in a significant proportion of cells, mitochondrial partitioning between the daughters though equitable, resulted in cells with very few mitochondria compared to the rest of the population. These cells that contained fewer mitochondria grew slower, and therefore would likely be out-competed by other cells. However, because the reduction in mitochondria resulted from altered MT dynamics, asymmetric cell division and thereby again daughter cells with fewer mitochondria, would persist in future division cycles. Dnm1Δ cells have previously been shown to have retarded growth rates (30), which could be partially attributed to the unequal partitioning of mitochondria following asymmetric cell division in a significant proportion of these cells.

## Conclusion

In conclusion, MT dynamics and mitochondrial form and attachment were found to be fine-tuned to be in a ‘Goldilocks zone’ in fission yeast whereby symmetric cell division could be achieved. Any deviation from this narrow range resulted in asymmetric cell division. Additionally, cellular homeostasis relied on the feedback between MTs and mitochondria, with the mitochondria dictating its own partitioning via changes in its form. In future, it will be interesting to understand the fate of cells that inherited fewer mitochondria, and if similar feedback mechanisms exist between the cytoskeleton and other intracellular compartments.

## Materials and Methods

### Strains and media

The fission yeast strains used in this study are listed in Table S1. All the strains were grown in yeast extract medium or Edinburgh minimal medium (EMM) with appropriate supplements at a temperature of 30°C (1).

### Construction of strains

Strain VA064 was constructed by transforming Dnm1Δ with pREP41-Dnm1 (Dnm1 untagged plasmid). Similarly, strain VA102 was constructed by crossing PT1650 (h+ cox4-GFP:leu1 ade6-M210 ura4-D18; see Table S1) with JCF4627 (h-ade6-M210 leu1-32 ura4-D18 his3-D1 hht1-mRFP-hygMX6; see Table S1) while strain VA103 was constructed by crossing VA077 (h-dnm1::kanr leu1-32ade-(ura+)cox4-GFP:leu1 ade6-M210 leu1-32 ura4-D18; see Table S1) with VA101 (h+ hht1-mRFP-hygMX6 cox4-GFP:leu1 ade6-M210 leu1-32 ura4-D18; see Table S1). Strain VA104 was constructed by crossing VA080 (h-mmb1Δ:Kanr cox4-GFP:leu2 mCherry-atb2:Hygr ade6-m210 leu1-32 ura4-d18; see Table S1) with VA101 (h+ hht1-mRFP-hygMX6 cox4-GFP:leu1 ade6-M210 leu1-32 ura4-D18; see Table S1). Strain VA110 was constructed by crossing VA109 (h+ dnm1Δ::kanr leu1-32ade-(ura+) ura4-Δ18 leu1::GFP-atb2+:ura4+; see Table S1) with JCF4627 (h-ade6-M210 leu1-32 ura4-D18 his3-D1 hht1-mRFP-hygMX6). Strain VA111 was constructed by crossing VA102 (h-hht1-mRFP-hygMX6 cox4-GFP:leu1 ade6-M210 leu1-32 ura4-D18; see Table S1) with MCI438 (h+ tea2d:his3 ade6 leu1-32 ura4-D18 his3-D1; see Table S1). Strain VA112 was constructed by crossing JCF4627 (h-ade6-M210 leu1-32 ura4-D18 his3-D1 hht1-mRFP-hygMX6; see Table S1) with VA106 (h+ ura4-Δ18 leu1::GFP-atb2+:ura4+; see Table S1). Strain VA113 was constructed by crossing VA112 (h+ hht1-mRFP-hygMX6 ura4-Δ18 leu1::GFP-atb2+:ura4+ ade6-M210 leu1-32 his3-D1; see Table S1) with VA078 (h+ mmb1Δ:Kanr; see Table S1).

### Plasmid transformation

Transformation of strains was carried out using the improved protocol for rapid transformation of fission yeast as described previously (16).

### Preparation of yeast for imaging

For imaging, fission yeast cells were grown overnight in a shaking incubator at 30°C. The following day, the cells were sub-cultured into fresh medium for 2 h at 30°C to achieve an optical density (OD) of 0.3-0.4 (mid-log phase). Following this, cells were washed once with distilled water and thrice with EMM. The cells were then allowed to adhere on lectin-coated (Sigma-Aldrich, catalog number L2380) 35-mm confocal dishes (SPL Life Sciences, cat. number 100350) for 20 min. Unattached cells were removed by washing with EMM.

### Live-cell imaging

Confocal microscopy was carried out in Fig. 1A, 4A, S1B and S2C using the InCell Analyzer-6000 (GE Healthcare) with a 60x air objective 0.95 numerical aperture (NA) objective fitted with an sCMOS camera. For GFP and RFP imaging, 488 and 561 nm laser lines and 525/20 and 605/52 nm bandpass emission filters, respectively, were used. Spinning disk confocal microscopy was carried out in Fig. 2A, 3A and S1A using the Eclipse Ti2-E (Nikon) with a 100× oil-immersion 1.49 NA objective fitted with an EM-CCD camera (iXon Ultra-897; Andor). For GFP and RFP imaging, 488 and 561 nm laser lines (Toptica) and 525/20 and 605/52 nm bandpass emission filters, respectively, were used.

Laser resonant scanning confocal microscopy was carried out in Fig. 5A, 5C, using the Nikon A1 with a 60x, water immersion, 1.2 NA objective fitted with GaAsP detectors. For GFP and RFP imaging, 488 and 561 nm laser lines and 525/50 and 595/50 nm bandpass emission filters, respectively with 405/488/561 dichroic, were used.

MT polymerisation, depolymerisation rates and MT pivoting in Fig. 2B was obtained by imaging Z-stacks (7 slices with step size 1 *μ*m) acquired every 3 s for 5 min. MT elongation times in Fig. 2D were imaged using Z-stacks (7 slices with step size 1 *μ*m) acquired every 7 s for 10 min. Short term nuclear dynamics in Fig. 3A were imaged using Z-stacks (7 slices with step size 1 *μ*m) acquired every 20 s for 20 min while long-term nuclear dynamics in Fig. S2C were imaged using Z-stacks (5 slices with step size 0.5 *μ*m) every 15 minutes for 12 hours. MT depolymerisation in Fig. S1A was observed in time-lapse movies containing Z-stacks (5 slices with step size 0.5 *μ*m) acquired every 12.5 s for 20 min. The growth rates of divided daughter cells in Fig. 5A and Fig. 5C was imaged with Z-stacks (13 slices with step size 0.5 *μ*m) every 7 min for 10 h and 14 min for 12 h respectively.

### Ultrastructure expansion microscopy

Ultrastructure expansion microscopy was performed as described in (17), with some modification to the cell fixation. Briefly, cells were grown in YES at 32°C for 36 h, followed by high pressure freezing. Cultures were concentrated onto nitrocellulose membranes by vacuum filtration and frozen in 200 μm aluminium carriers at an ABRA HPM010. Freeze substitution was performed at -90°C in acetone (Sigma, cet. No. 24201M) and gradually warmed to room temperature at 5°C/h. Cells were subsequently rehydrated by successive washes with EtOH containing increasing amounts of H_2_O (0%, 0%, 5%, 5%, 25%, 50%, 100%, five minutes each) and stored until further use in PBS at 4°C (https://doi.org/10.1038/s41592021-01356-4). For cell wall digestions, fixed cells were rinsed once in PEM buffer (100 mM PIPES, 1 mM EGTA and MgSO4, pH 6.9) and 2x in PEM containing 1.2 M sorbitol (PEMS) before incubating them in 2.5 mg/mL Zymolyase 20T (Roth, cat.no. 9324.3) in PEMS at 37°C with agitation for 45 min. Cell wall digestion was confirmed with calcofluor white staining, and cells were then washed 3x in PEMS buffer. The resulting cell suspension was loaded onto a 12mm lysine-coated coverslip and processed for expansion.

The coverslips now containing fixed spheroplasts were incubated in protein crosslinking prevention solution (2% acrylamide (AA, Sigma, cat.no. A4058) / 1.4% formaldehyde (Sigma, cat.no. F8775) in PBS) for 3 to 5h at 37°C. To the monomer solution (19% (wt/wt) sodium acrylate (Sigma, cat.no. 408220), 10% (wt/wt) AA, 0.1% (wt/wt) N,N’-methylenbisacrylamide (Sigma, cat.no. M1533) in 1X PBS), ammonium persulphate (ThermoFisher, cat.no. 17874) and tetramethylethylendiamine (ThermoFisher,cat.no. 17919)) were added to a final concentration of 0.5% each and the gelation was performed in a pre-cooled humid chamber on ice for 5 min and at 37°C for 1 h. The coverslips were then incubated in denaturation buffer (50 mM Tris pH 9, 200 mM NaCl, and 200 mM SDS in water, pH 9) with agitation for 15 min at room temperature. The formed gels were then transferred to Eppendorf tubes containing denaturation buffer and incubated for 90 min at 95°C without agitation. Gels were expanded with 3x baths of ddH2O for 30 min at RT. After full expansion of the gel, the diameter of the gel was measured and proceed for immunostaining with NHS-ester diluted (at 2μg/mL in PBS over night at 4°C) for visualisation of the general organisation of the cell (including mitochondria and the nucleus), and YL1/2 rat anti-*α*tubulin antibody (gift from Gislene Pereira) for visualisation of MTs. Expanded cells were then imaged using a spinning disk confocal microscope (Olympus IXplore SpinSR, with 0.95NA 40x air objective; Z-stacks spanning the entire cells were taken with 0.3 μm step size.

### Image and data analysis

Images were analysed using Fiji/ImageJ (31). Interphase cells that were used in our analyses had a mean length of 10 *μ*m. This mean length corresponds to cells in early-mid G2 phase in *S. pombe* (32).

For analysis of the length of MT that was attached to mitochondria in Fig. 1A, the colocalisation of MTs with mitochondria in WT and Dnm1Δ in ultrastructure expanded cells was measured through the Z-stack containing the entire cell, and summed for each cell. The summed values were then normalised to the mean of the WT.

The MT polymerisation and depolymerisation rates were obtained by measuring the angle of the slopes (*θ*) from kymographs generated by drawing a line along a growing or shrinking MT and using the following formula:

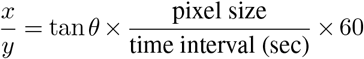

Where:

x: is the MT length in μm

y: is the time in min

The MT elongation time was calculated from the kymograph by measuring the time from the onset of polymerisation to a catastrophe event. The rate of catastrophe was obtained from the reciprocal of the mean elongation time. The nuclear dynamics were obtained by thresholding the nucleus from time-lapse videos in ImageJ to obtain the nuclear centroid, and drawing an ROI around the cell perimeter to get the cell centroid. Then the euclidian distance between the 2 centroids was then calculated.

The nuclear velocity in Fig. S2D by measuring the euclidean distance between the nuclear positions in successive frames. MT pivoting was measured as the difference in the angle of the MT from one frame to another.

In Fig. 4B, mitochondrial intensities in daughter cells were normalised to total mitochondrial intensity of the mother, and similarly, area of the daughter cells was normalised to the total area of the mother cell just prior to division.

In Fig. 5A and S2E, the cell area is measured in each frame from the first to the last frame and all the cell areas are normalised to the cell area in the first frame.

In Fig. 5B and S2F, the normalised mitochondrial intensity represents the mitochondrial intensities of the daughter cells at birth divided by the mitochondrial intensity of the mother cell. The growth rate represents the rate of change of cell area between the first and last frames of the timelapse images. Only cells with <20% asymmetry were used for quantification.

### Statistics and plotting

Data were checked for normality using the chi2gof function in Matlab. Then, to test the statistical significance of the difference between distributions we used ordinary one-way ANOVA or student’s T-test for parametric data and Kruskal-Wallis test or Mann-Whitney test for non-parametric data. Equality of variance was compared using Levene’s test. All plots were generated using Matlab (Mathworks Corp.). The figures were organised and prepared in Illustrator.

## Acknowledgements

We thank Ananya Rajagopal for help with construction of strains; the Katharina Gaus Light Microscopy Facility, UNSW; the High Content Imaging Facility, Centre for BioSystems Science and Engineering, Indian Institute of Science for the use of the InCell 6000 and spinning disk confocal microscopes; G. Redpath, N. Ul Fatima, A. Badrinarayanan, N. Dua, M. Rao and I. Jain for comments on the manuscript; P. Delivani (Max Planck Institute of Molecular Cell Biology and Genetics, Dresden, Germany), M. Takaine (Gunma University, Gunma, Japan), I. Tolić (Ruder Bošković Institute, Zagreb, Croatia), P. Tran (University of Pennsylvania, Philadelphia, PA), T.D. Fox (Cornell University, Ithaca, NY), and National BioResource Project Japan for yeast strains and constructs.F. Mikus and G. Dey acknowledge the European Molecular Biology Laboratory for support.

