## Supplementary Information for "Microtubule-mitochondrial attachment facilitates cell division symmetry and proper mitochondrial partitioning in fission yeast"

This document contains 2 Supplementary Figures, one Supplementary Table, and Supplementary Video Captions.

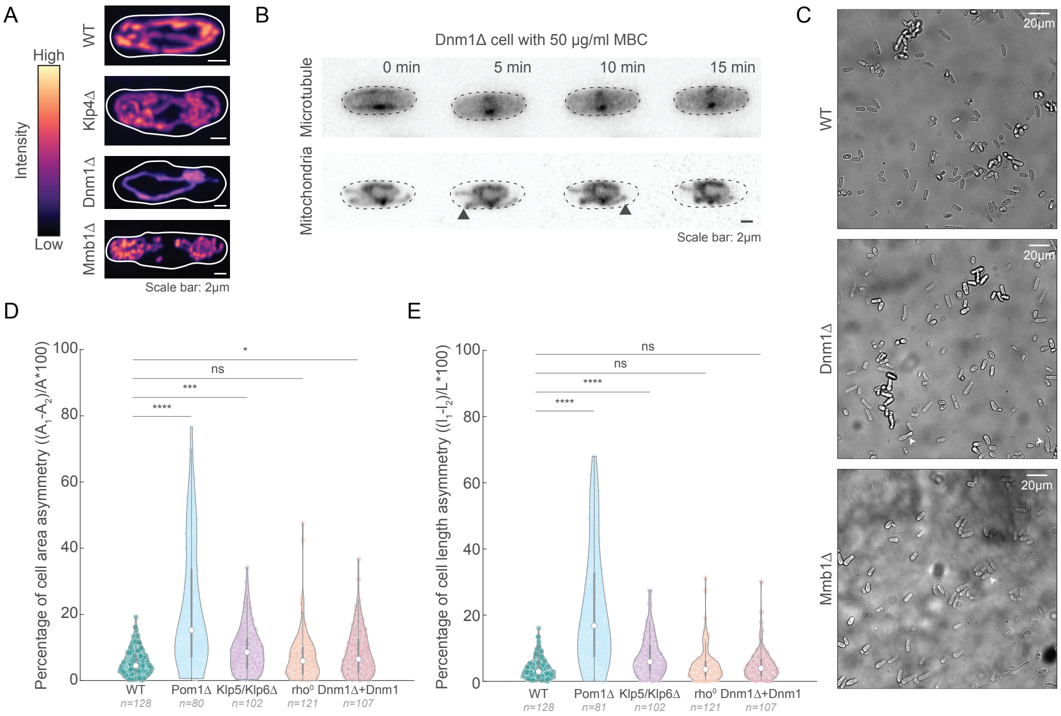


**Fig. S1. Pom1△ and Klp5/6△ exhibit asymmetric cell division. A**, Maximum intensity-projected images of mitochondria (top to bottom) in WT, Klp4△, Dnm1△ and Mmb1△ cells (strains VA102, VA111, VA103, VA104, see Table S1), represented in the intensity map located to the left of the images. **B**, Montage of maximum intensity-projected images of MTs and mitochondria in a representative Dnm1△ cell (strain VA069,see Table S1) treated with 50$\mu$g/ml MBC. The change in position of mitochondria is indicated with the grey arrowhead at times 5 and 10 min. We noticed this phenomenon in 100% of the cells observed (n=50). **C**, Representative bright-field images of the whole field of view of WT (top), Dnm1△ (middle) and Mmb1△ (bottom) cells. The white arrowheads point to examples of asymmetrically dividing cells. **D**, Plot of asymmetry in cell areas between the daughter cells in WT, Pom1△, Klp5/6△, *rho*$^0$ and Dnm1△+Dnm1 plasmid cells (strains L972, FY7143, KI001, FY31851, G3B, PHP14 and VA064, see Table S1). **E**, Plot of asymmetry in cell lengths between the daughter cells in WT, Pom1△, Klp5/6△, *rho*$^0$ and Dnm1△+Dnm1 plasmid (strains L972, FY7143, KI001, FY31851, G3B, PHP14 and VA064, see Table S1).

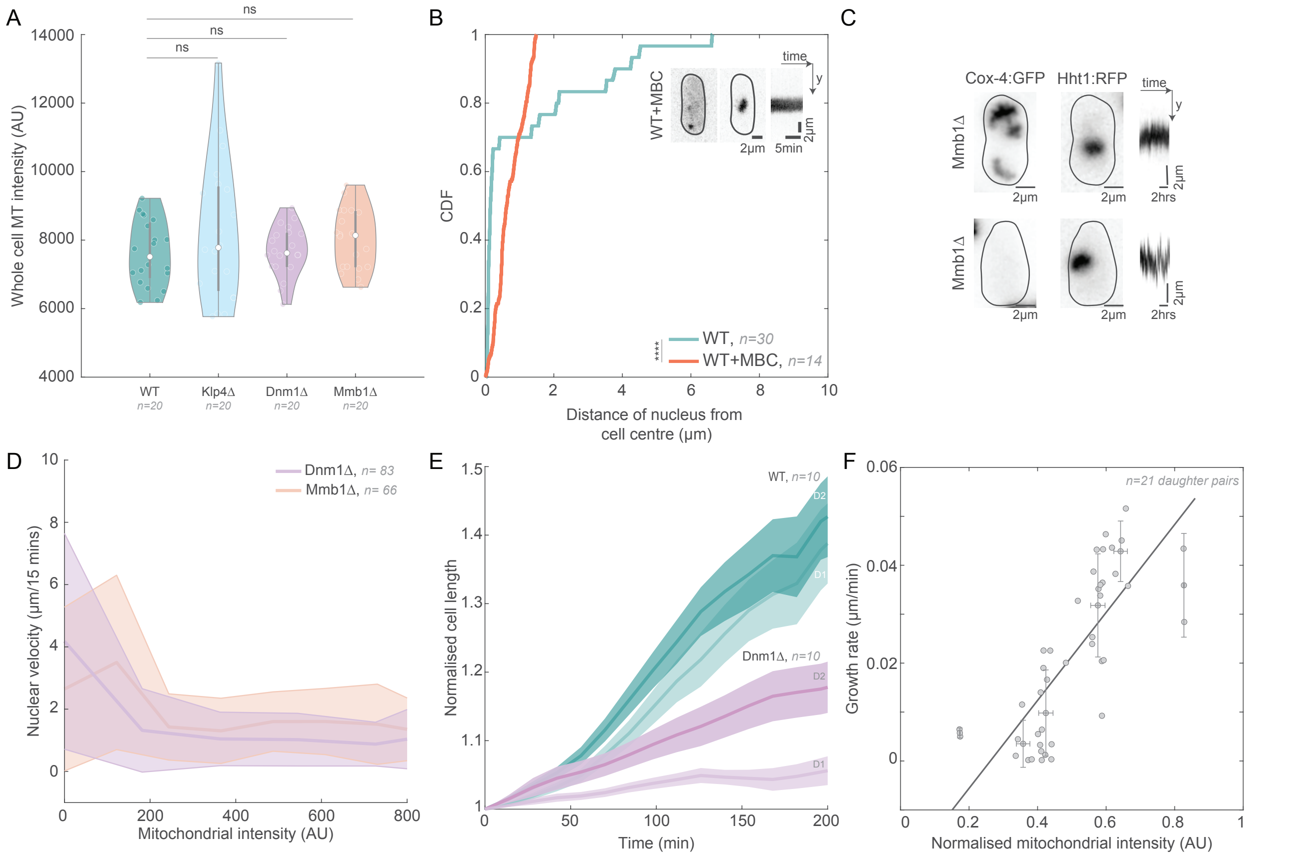


**Fig. S2. The nucleus is more dynamic when mitochondria are present in low numbers.** **A**, Plot of summed intensity of the MT in whole WT, Klp4Δ, Dnm1Δ and Mmb1Δ cells (strains VA112, G5B, VA110, and VA113, see Table S1). 'ns' indicates no significant difference using ordinary one-way ANOVA for parametric data. **B**, CDF of the distance of the nucleus from the cell centre for each time point of the time-lapse videos of nuclei in WT cells and WT cells with 50$\mu$g/ml MBC. Note that the WT data has been reproduced from Fig. 4B. Maximum intensity-projected images (top right) of depolymerised MTs and histone in WT+MBC cells (strain VA102, see Table S1). The asterisks (****) represent p<$10^{-4}$, Mann-Whitney Test for non-parametric data. **C**, Maximum intensity-projected images of mitochondria (left) and nucleus (centre) from the first frame of time-lapse videos of representative Mmb1Δ cells (strain VA104, see Table S1). Kymograph (right) depicts nuclear movement from the first to last frame. **D**, Plot of nuclear velocity measured every 15 minutes versus the summed intensity of the mitochondria in Dnm1Δ and Mmb1Δ (strains VA103 and VA104, see Table S1). **E**, Plot of change in cell length vs. time of smaller (D1) and larger (D2) cells normalised to the first time frame upon division of the mother cell. 10 D1-D2 pairs were analysed for WT and Dnm1Δ cells (strains VA102 and VA103, see Table S1). **E**, Plot of growth rate vs. mitochondrial intensities in 21 Mmb1Δ daughter cell pairs that underwent symmetric cell division (<20% asymmetry between daughters). The black line is a weighted linear fit (of the form $y = mx + c$), and yielded $R^2$ =0.77.

### Supplementary Table

| Name | Genotype | Source |
|---|---|---|
| L972 | h- WT | Iva Tolic' |
| FY7143 | h- ura4-D18 leu1-32 ade6-M216 his7-366 | YGRC, Japan |
| KI001 | h+ sid4-GFP::kan r kan r -nmtP3-GFP-atb2+ nmt1-pCOX4-RFP::leu1+ ura4-D18 ade6-M210 | Iana Kalinina |
| G5B | h- klp4::kanr nmt1-GFP-atb2 leu ura | Rafael Carazo Salas, UK |
| FY31851 | h- leu1-32 CRIB:GFP(ura+) pom1::ura4+ rga4:RFP(kanMX6) | YGRC, Japan |
| Dnm1Δ | h- dnm1::kanr leu1-32ade- | Yannick Gachet, Toulouse |
| PT2244 | h+ mmb1Δ:Kanr cox4-GFP:leu2 mCherry- atb2:Hygr ade6-m210 leu1-32 ura4-d18 | Phong Tran, USA |
| PT1650 | h+ cox4-GFP:leu1 ade6-M210 ura4-D18 | Phong Tran, USA |
| JCF4627 | h- ade6-M210 leu1-32 ura4-D18 his3-D1 hht1-mRFP-hygMX6 | Julie Cooper Lab |
| MCI438 | h+ tea2d:his3 ade6 leu1-32 ura4-D18 his3-D1 | Iva Tolić, Croatia |
| PHP 14 | h- ade6-M216, leul-32, ptp-1, [rho0] | Thomas D. Fox |
| VA064 | h- pREP41-Dnm1(leu+) dnm1::kanr leu1-32 ade-(ura+) | This study |
| VA069 | h- pREP1-atb2:GFP(leu+) dnm1::kanr leu1-32 ade-(ura+)) | This study |
| VA077 | h- dnm1::kanr leu1-32ade-(ura+) cox4-GFP:leu1 ade6-M210 leu1-32 ura4-D18 | This study |
| VA078 | h+ mmb1Δ:Kanr | This study |
| VA080 | h- mmb1Δ:Kanr cox4-GFP:leu2 mCherry-atb2:Hygr ade6-m210 leu1-32 ura4-d18 | This study |
| VA101 | h+ hht1-mRFP-hygMX6 cox4-GFP:leu1 ade6-M210 leu1-32 ura4-D18 | This study |
| VA102 | h- hht1-mRFP-hygMX6 cox4-GFP:leu1 ade6-M210 leu1-32 ura4-D18 | This study |
| VA103 | h- hht1-mRFP-hygMX6 dnm1::kanr cox4-GFP:leu1 ade6-M210 ura4-D18 | This study |
| VA104 | h- hht1-mRFP-hygMX6 mmb1Δ:Kanr cox4-GFP:leu1 ade6-M210 ura4-D18 | This study |
| VA106 | h+ ura4-Δ18 leu1::GFP-atb2+:ura4+ | This study |
| VA109 | h+ dnm1Δ::kanr leu1-32ade-(ura+) ura4-Δ18 leu1::GFP-atb2+:ura4+ | This study |
| VA110 | h- dnm1Δ::kanr leu1-32ade-(ura+) ura4-Δ18 leu1::GFP-atb2+:ura4+ hht1-mRFP-hygMX6 | This study |
| VA111 | h- tea2d:his3 hht1-mRFP-hygMX6 cox4-GFP:leu1 ade6-M210 ura4-D18 | This study |
| VA112 | h+ hht1-mRFP-hygMX6 ura4-Δ18 leu1::GFP-atb2+:ura4+ ade6-M210 leu1-32 his3-D1 | This study |
| VA113 | h+ mmb1Δ:Kanr hht1-mRFP-hygMX6 ura4-Δ18 leu1::GFP-atb2+:ura4+ ade6-M210 leu1-32 his3-D1 | This study |
| pREP41-Dnm1 (plasmid) | Dnm1 (untagged) | Isabelle Jourdain, UK |

#### Supplementary Video Captions

**Video S1. Mitochondria in a Dnm1Δ cell retracts upon addition of MBC**
A representative Dnm1Δ cell stained with 200nM MitoTracker Orange and expressing atb2:GFP (strain VA069, see Table S1) imaged using confocal microscopy every 12.5 s. Scale bar: 2$\mu$m, Time is indicated in mm:ss.

**Video S2. Ultrastructure expansion microscopy reveals increased attachment of mitochondria to MTs in Dnm1Δ cells**
3D projections of representative WT, Dnm1Δ and Mmb1Δ cells stained for the MTs (green) and mitochondria/nucleus (strains L972, Dnm1Δ and VA073, see Table S1) visualised using a spinning disk confocal microscope following ultrastructure expansion.

**Video S3. The nucleus is more dynamic in Dnm1Δ and Mmb1Δ cells**
WT, Klp4Δ, Dnm1Δ and Mmb1Δ cells expressing Hht1-mRFP (strain VA102, VA103, VA104 and VA111, see Table S1) imaged using spinning disk microscopy every 20 s. Scale bar: 2$\mu$m, Time is indicated in mm:ss.

**Video S4. Nuclear dynamics is increased when mitochondria are present in low amounts**
An Mmb1Δ cell expressing Cox4:GFP and Hht1:mRFP (strain VA104, see Table S1) imaged using confocal microsocpy every 15 min. Scale bar: 2$\mu$m, Time is indicated in hh:mm.

**Video S5. The larger daughter cell grows faster than the smaller one after the mother cell divides asymmetrically**
A WT (top) and Dnm1Δ (bottom) cell expressing expressing Cox4:GFP and Hht:mRFP (strain VA102 and VA103 respectively, see Table S1) imaged using confocal microscopy every 7 min. Scale bar: 2$\mu$m.

**Video S6. Growth rate of cells scales with the amount of mitochondria inherited at birth**
An Mmb1Δ cell expressing Cox4:GFP and Hht1:mRFP (strain VA104, see Table S1) imaged using confocal microscopy every 14 min. Scale bar: 2$\mu$m.
